# BubR1 and Mad2 regulate adult midgut remodeling in *Drosophila* diapause

**DOI:** 10.1101/2025.01.26.634899

**Authors:** Yuya Adachi, Hiroki Nagai, Tenki Nakasugi, Miyu Nagao, Hiroko Nakatani, Takako Fujichika, Aya Takahashi, Masayuki Miura, Yu-ichiro Nakajima

## Abstract

Diapause is a survival strategy in which growth and aging are temporarily suspended, enabling animals to withstand unfavorable environments. Various insects, including the fruit fly *Drosophila*, enter reproductive diapause, or dormancy, in response to colder temperatures and/or shorter day lengths. During reproductive diapause, ovarian development halts, and non-reproductive organs also undergo remodeling at both morphological and metabolic levels: however, the mechanisms underlying this remodeling and its physiological impact remain largely unclear. Here, we show that the *Drosophila* adult midgut undergoes extensive remodeling in diapause, marked by a sustained suspension of growth due to the cell cycle arrest of intestinal stem cells (ISCs) that revert to normal upon returning to recovery conditions. During dormancy, BubR1 and Mad2, key regulators of mitosis, are highly expressed and localized in the cytoplasm of ISCs rather than at the kinetochore, and both BubR1 and Mad2 are essential for diapause-specific midgut remodeling. Furthermore, disruption of midgut growth arrest during diapause reduces the resistance to starvation in adult flies. Together, our findings identify a novel role for BubR1 and Mad2 in ISCs–promoting proper midgut remodeling during dormancy–and highlight the importance of this process for survival under adverse environments.

## Introduction

Wild animals have developed a wide array of survival strategies to cope with harsh environmental conditions such as temperature fluctuations, drought, and food shortages. Among these strategies, dormancy programs (e.g. hibernation, torpor, and diapause) are critical adaptations that involve a reduction in metabolic rate and body temperature, which conserves energy, extends lifespan, and increases the probability of survival for the species (Easwaran & Montell, 2023; Staples, 2016; Wilsterman et al., 2021). By temporarily halting growth and delaying organ development under stressful conditions such as starvation and low temperatures, dormancy helps reduce the energy expenditure required for growth and reproduction, and this synchronization of physiological processes with environmental conditions enhances the organism’s ability to survive (Denlinger, 2022; Schiesari & O’Connor, 2013). Understanding the underlying mechanisms of dormancy may thus provide valuable insights into the regulation of stress resistance, aging, and reproduction.

Dormancy programs typically comprise three distinct phases: a preparatory phase in which environmental cues initiate dormancy; a dormancy phase characterized by reduced metabolism and activity; and a recovery phase wherein normal growth and activity resume (Gray et al., 1991; Harsimran Kaur Gill et al., 2017). To enter and exit dormancy at the appropriate time, animals must be able to successfully detect environmental conditions and coordinate internal physiological processes. As such, dormancy regulation generally requires central control mechanisms involving the brain and endocrine systems (Helfrich-Förster, 2024; Karp, 2021). While hormonal and neural regulation play pivotal roles in directing these processes, peripheral organs and tissues must also undergo significant remodeling during dormancy and subsequent reversion during recovery. For instance, during hibernation in certain mammals (e.g. thirteen-lined ground squirrels, Syrian hamsters) and diapause in insects, structural changes are observed in respiratory and digestive organs as well as in muscles and adipose tissues (Kim et al., 2006; Kim & Denlinger, 2009; Kurtz et al., 2021). Despite these substantial transformations and their vital importance, the mechanisms by which cellular compositions within individual organs and tissues are precisely coordinated to adjust dormancy programs remain largely unknown.

Animal species enter dormancy at different life stages, and in adult insects, dormancy is often referred to as reproductive diapause, a state that suppresses reproductive organ development and is associated with reduced locomotor activity, lower metabolism, increased stress resistance, and extended lifespan(Denlinger, 2022; Tatar & Yin, 2001). In species like *Drosophila*, reproductive diapause is triggered by low temperatures and/or short-day conditions (Hara & Yamamoto, 2022; Lirakis et al., 2018; Ojima et al., 2018; Saunders et al., 1989). As with other diapausing insects, the reduction of juvenile hormone and insulin signaling are critical for maintaining reproductive diapause in *Drosophila* (Easwaran et al., 2022; Saunders et al., 1990; Schiesari et al., 2016). Recent studies have used *D. melanogaster* to explore the genetic basis of dormancy’s complex biology and have resulted in the identification of circadian neurons that regulates the endocrine organs, the mechanism of germline maintenance, and sleeping behaviors (Easwaran et al., 2022; Kurogi et al., 2023; Meiselman et al., 2022; Meyerhof et al., 2024; Schiesari et al., 2016). Interestingly, similar to dormancy programs in other animals, somatic non-reproductive organs such as the midgut undergo structural changes during reproductive diapause in *Drosophila* (Kubrak et al., 2014); however, the mechanism by which the dormant state regulates organ remodeling and its physiological impacts remain unclear.

The *Drosophila* adult midgut, composed of a monolayered epithelium surrounded by visceral muscle, is home to resident intestinal stem cells (ISCs) that self-renew and give rise to committed progenitors, enteroblasts (EBs) and enteroendocrine progenitor cells (EEPs); these progenitors differentiate into absorptive enterocytes (ECs) and secretory enteroendocrine cells (EEs), respectively (Hung et al., 2020; Miguel-Aliaga et al., 2018; see also Fig. 2A, 2B). To achieve adaptive responses to environmental changes at the organ level, the ISC lineage exhibits characteristic behaviors that enable proper tissue homeostasis and plasticity (Nagai et al., 2022; O’Brien et al., 2011). Furthermore, the midgut plays a central role in regulating organism-level physiology, influencing appetite, fecundity, immunity, and lifespan (Malita et al., 2022; White et al., 2021; Yoshinari et al., 2024; Zhou et al., 2020). With its relatively simple structure and conserved cell lineages, the *Drosophila* adult midgut serves an ideal system for dissecting the mechanisms of organ remodeling associated with environmental challenges like dormancy.

Here, we investigate the mechanistic underpinnings of organ remodeling during diapause, using the *Drosophila* adult midgut as a model. Our results show that intestinal growth halts during dormancy and resumes during recovery, accompanied by changes in structural and metabolic gene expression. We identify mitotic regulators BubR1 and Mad2, which are highly expressed in the cytoplasm of ISCs, as key players in midgut remodeling during diapause, and show that disruption of these genes leads to abnormal gut size and a reduced capacity for starvation resistance. These findings suggest novel roles of BubR1 and Mad2 in controlling ISCs dynamics and contributing to midgut remodeling in *Drosophila* diapause.

## Results

### The adult midgut of *D. melanogaster* undergoes remodeling in response to diapause and recovery

In adult females of *D. melanogaster*, reproductive diapause can be induced by exposing newly eclosed flies to cold temperatures (below 12°C), halting ovarian development at an early stage (Anduaga et al., 2018). Following previous studies (Easwaran et al., 2022; Kubrak et al., 2014; Saunders et al., 1989), we kept newly eclosed flies at short-day photoperiods (10L:14D) and 10°C while non-diapause control flies were kept at 12L:12D and 18°C (Fig. 1A). Under this suboptimal condition, females were confirmed to have entered reproductive dormancy as no yolk accumulation was observed in the ovaries (Fig. S1A). To assess changes in midgut size during diapause, we measured the length and area of the midgut in the wild-type fly (Canton-S) under diapause and non-diapause conditions for 1 to 5 weeks. Consistent with previous findings (Kubrak et al., 2014), we confirmed that midgut length remained shorter during diapause compared to non-diapause conditions (Fig. 1B, 1C), while becoming similar to that immediately after eclosion (Fig. 1C). After one week of recovery from diapause, the midgut significantly increased in size (Fig. 1B, 1D, Fig. S1B, S1C) and fully recovered by day 4 (Fig. S1D). Of note, after 5 weeks of diapause, no significant midgut growth occurred upon recovery (Fig. 1D), suggesting the existence of a time limit on the midgut’s ability to regrow. These findings together indicate that the midgut halts growth during diapause but remains capable of rapid regrowth upon recovery.

**Figure 1.**
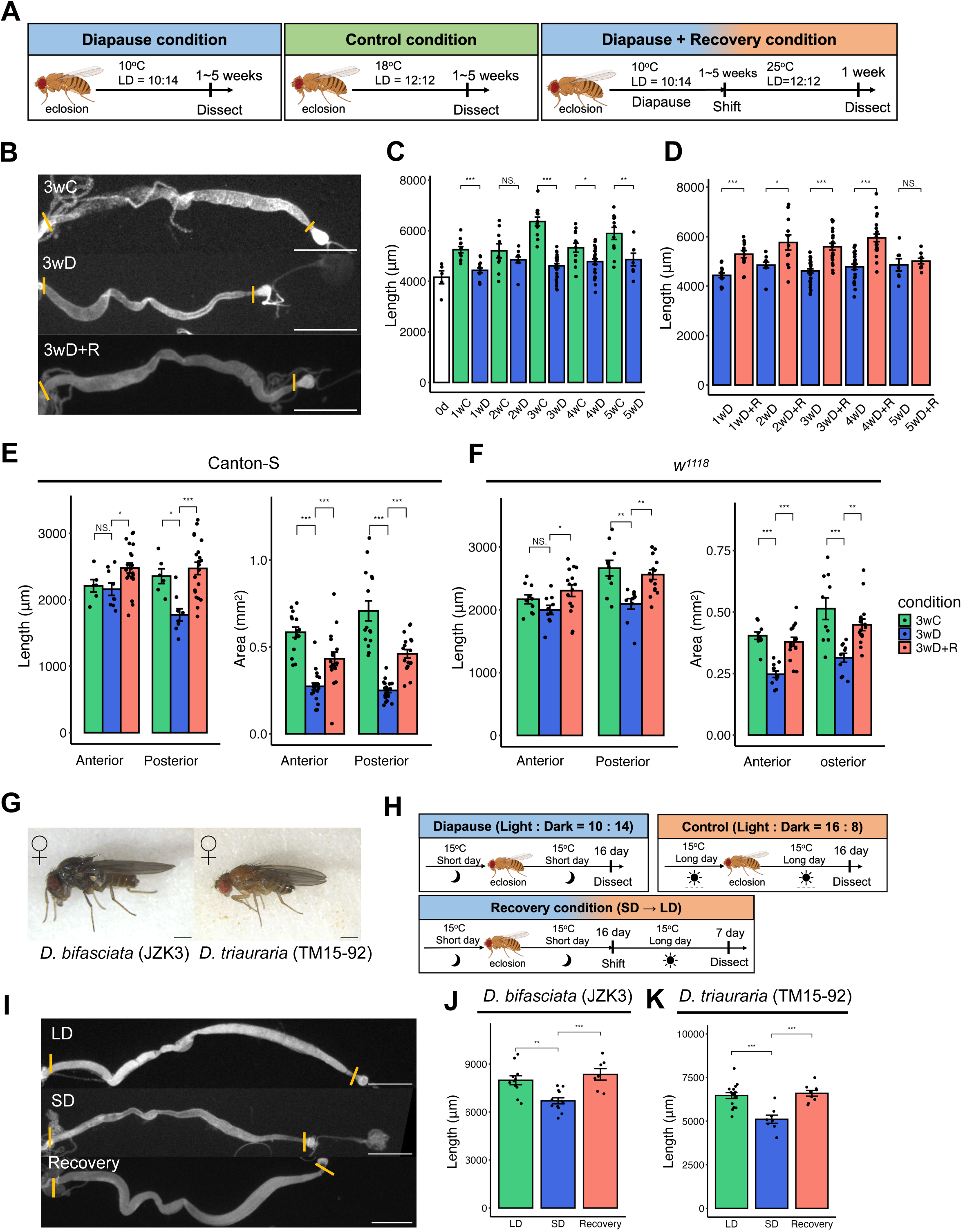
The adult midgut of *Drosophila* undergoes remodeling in response to diapause and recovery. (A) Schematic of diapause and recovery in *D. melanogaster*. LD = Light: Dark. (B) Representative images of the midgut under the 3 weeks diapause (3wD) and recovery (3wD+R) condition. The two orange lines indicate anterior and posterior end of the midgut. Scale bars: 1 mm. (C, D) Quantification of midgut length of wild-type flies (Canton-S) under diapause and control conditions (C) and their recovery conditions (D). *n=*10-15 midguts for each condition. The same diapause samples are quantified in (C) and (D). (E) Quantification of midgut length and area in the anterior and posterior region. Canton-S was analyzed. *n=*6 (3wC), 9 (3wD), and 21 (3wD+R) midguts for length measurements and *n=*15 (3wC), 20 (3wD), 17(3wD+R) midguts for area measurements. (F) Quantification of midgut length and area in the anterior and posterior region. *w^1118^*was analyzed. *n*=10 (3wC), 10 (3wD), 15 (3wD+R) midguts for length measurements and *n*=10 (3wC), 11 (3wD), 15 (3wD+R) midguts for area measurements. (G) Female adult of *D. bifasciata* (JZK3) and *D. triauraria* (TM15-92). Scale bars: 1 mm. (H) Schematic of diapause and recovery conditions in *D. bifasciata* and *D. triauraria.* Light: Dark= 16: 8 (hours) for long day (LD) condition, 10: 14 (hours) for short day (SD) condition. (I) Representative images of the midgut in *D. bifasciata* (JZK3) under control (LD), diapause (SD), and recovery conditions. Scale bars: 1 mm. (J, K) Quantification of midgut length in *D. bifasciata* (J) and *D. triauraria* (K). *n*=7-12 midguts for each condition.

The *Drosophila* adult midgut consists of anterior (R1-R2), middle (R3), and posterior regions (R4-R5) (Buchon et al., 2013; Marianes & Spradling, 2013). Since these regions respond differently to environmental changes due to the divergent nature of ISCs and their progenies (Li & Jasper, 2016; Mattila et al., 2024; Nagai et al., 2023; O’Brien et al., 2011), we analyzed midgut size changes in the anterior and posterior regions under the 3-week diapause condition where the difference between diapause and non-diapause midgut size is greatest (Fig. 1C). We found that the posterior midgut differed more significantly across conditions (Fig. 1E) than the anterior midgut. Similar region-specific gut resizing during diapause and recovery was also observed in the *w^1118^* strain (Fig. 1F), suggesting that the posterior region is preferentially affected during dormancy. It has been reported that reproductive development is also suppressed in *Drosophila* adult males under the same conditions females enter diapause (Kubrak et al., 2016). To assess diapause’s effect on the midgut across sexes, we measured midgut size in males under diapause and non-diapause conditions but ultimately found no significant differences between the two (Fig. S1G). These observations suggest that midgut remodeling in diapause is a sexually dimorphic trait, appearing only in females.

To determine whether the observed midgut remodeling is specific to diapause or is rather a general response to cold-induced stress, we exposed female flies to the cold temperature conditions outlined above after the midgut had matured and ensuring no diapause was induced (Ojima et al., 2018). Comparing the midgut size after 3 weeks under cold and diapause conditions, we found that midgut size was larger under cold conditions (Fig. S1E, S1F). Furthermore, midgut growth was not observed when flies were transferred from the cold condition to recovery, contrary to the transition observed during diapause recovery (Fig. S1F). Altogether, these results suggest that midgut remodeling is a diapause-specific response in adult females in which midgut growth is halted in a regrowth-ready state. We term this midgut remodeling during diapause: “dormancy-dependent midgut remodeling” (DDMR).

### Midgut remodeling as a common response in *Drosophila* species

While the model organism *D. melanogaster* has recently been used as a genetic model to study numerous aspects of diapause (Easwaran et al., 2022; Easwaran & Montell, 2024; Meiselman et al., 2022; Ojima et al., 2018), it is still unclear whether this species naturally enters diapause in the wild (Lirakis et al., 2018). However, it has been established that some species of the genus *Drosophila* undergo a programmed state of diapause in response to environmental changes like short-day lengths that enables them to survive through the winter (Kimura, 1990; Lakovaara et al., 1972).

To explore whether DDMR is a general organ-level response during reproductive dormancy, we examined two other *Drosophila* species: *D. bifasciata* and *D. triauraria* (Fig. 1G), which inhabit northern regions and have been shown to halt ovarian development in short-day seasons (Watabe, 1979; Yamada & Yamamoto, 2011). We confirmed that these species exhibit reproductive diapause under short-day conditions (SD, 10L: 14D), while such diapause was not induced in long-day conditions (LD, 16L: 8D) at the same moderate temperature (15°C) (Fig. 1H, S1H). In SD, midgut size was significantly smaller compared to LD, and midgut regrowth was observed during recovery (Fig. 1I-1K). Notably, this midgut growth arrest was not observed in non-diapause strains of *D. triauraria* that originate from southern regions (Yamada & Yamamoto, 2011) (Fig. S1I, S1J). These results support the notion that DDMR in *D. melanogaster* represents a general organ-level response triggered during diapause in the genus *Drosophila*.

### ISCs maintain cell cycle arrest and inhibition of mitosis during diapause

In the adult midgut epithelium of *Drosophila*, ISCs serve as resident stem cells, maintaining the self-renewing population and producing progenitors (EBs and EEPs) that further differentiate into absorptive ECs and secretory EEs, respectively (Fig. 2A, 2B; Hung et al., 2020; Miguel-Aliaga et al., 2018). Previous studies have shown that alterations in cell number via ISC expansion and reduction contribute to the substantial midgut size changes that correspond with environmental fluctuations (Christensen et al., 2024; Nagai et al., 2022; Stojanović et al., 2022). For example, after mating, juvenile hormones stimulate ISC proliferation, leading to an increase in midgut size (Reiff et al., 2015). Similarly, upon refeeding following starvation, ISC symmetric division is promoted in the posterior midgut (O’Brien et al., 2011), resulting in ISC expansion and a subsequent increase in total cell number as well as midgut size.

**Figure 2.**
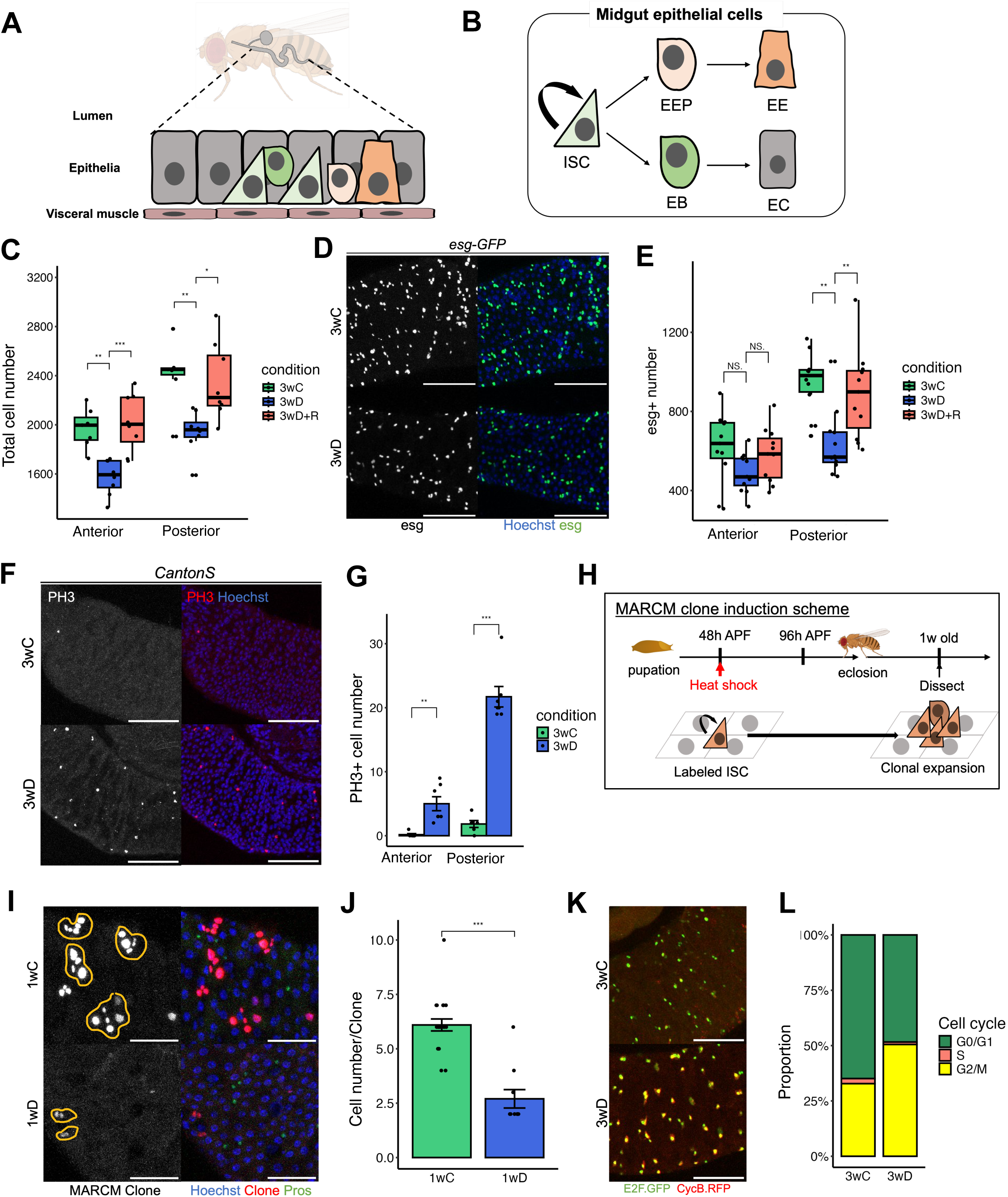
ISCs maintain cell cycle arrest and mitotic inhibition throughout diapause. (A, B) Schematics of the structure (A) and cell types (B) of the adult midgut of *D. melanogaster*. An intestinal stem cell (ISC) generates an enteroblast (EB) or an enteroendocrine progenitor (EEP), which differentiates into an enterocyte (EC) or an enteroendocrine cell (EE), respectively. (C) Total cell number in the adult midgut of wild-type flies under 3 weeks control (3wC), 3 weeks diapause (3wD), and recovery after 3 weeks diapause (3wD+R). *n*=7 (3wC), 8 (3wD), 8 (3wD+R) midguts. (D) Representative images of *esg-GFP^+^* cells in the posterior midgut under 3wC and 3wD conditions. Scale bars: 100 μm. (E) Quantification of *esg-GFP^+^*cell number. *n*=11 (3wC), 12 (3wD), 10 (3wD+R) midguts. (F) Representative images of anti-PH3 staining in 3wC and 3wD wild-type flies. Scale bars: 100 μm. (G) PH3+ cell number in the anterior and posterior regions of 3wC and 3wD flies. *n*=6 (3wC), 8 (3wD) midguts. (H) Schematics of MARCM clone induction. ISC was labeled before eclosion, and clonal expansion was examined at 1 week after eclosion. (I) Representative images of MARCM clones in 1wC and 1wD conditions. A single clone derived from an ISC is enclosed within the orange lines. Scale bars: 50 μm. (J) Quantification of clone size. *n*=21 (1wC),10 (1wD) clones. (K, L) Representative images of the Fly-Fucci signals in the progenitor (esg+) cells (K). Scale bars: 100 μm. The proportion of G2/M (GFP+ and RFP+ cells) is higher under the 3wD condition (L). *n*=7 (3wC), 5 (3wD) midguts.

To investigate whether changes in cell number contribute to midgut remodeling in diapause, we first quantified the midgut cell abundance following 3 weeks of diapause. We observed a significant reduction in total cell number during diapause compared to non-diapause conditions (Fig. 2C). We then examined the number of ISC/EB lineage using the protein trap line *esg-GFP.* The number of *esg*+ cells, along with their relative proportion, was markedly lower during diapause, especially in the posterior midgut (Fig. 2D, 2E, Fig. S2A). Consistent with this observation, the ISC marker *Delta* (*Dl*) and the EB marker *Su(H)* exhibited reduced expression in the diapausing midgut (Fig. S2B-S2F). Moreover, *esg*+ cell number dramatically increased during recovery from diapause, coinciding with midgut regrowth (Fig. 2E). These findings suggest the possibility that ISCs remain in a non-proliferative state during diapause and resume proliferation upon recovery.

To test this possibility, we investigated ISC mitotic activity using the anti-phospho-Histone H3 (PH3) immunostaining, a marker for mitotic cells. Unexpectedly, we found that the number of ISCs in the M phase significantly increased in both the anterior and posterior midgut during diapause (Fig. 2F, 2G). Despite the reduced number of ISCs (Fig. 2E, S2B, S2C), the increase in mitotic cells suggests that ISCs may arrest the cell cycle in the M phase or, alternatively, experience prolonged M phase during diapause.

In order to directly test ISC proliferative capacity, we conducted clonal expansion assays using the mosaic analysis with a repressible cell marker (MARCM) system (Fig. 2H) and found that clonal size was significantly reduced during diapause, with fewer ISCs present in the clones (Fig. 2I, 2J). This mitotic arrest was not observed during recovery from diapause, with the number of mitotic cells returning to non-diapause levels (Fig. S2G).

To further examine the ISC lineage cell cycle state, we employed the Fly-FUCCI system (Zielke et al., 2014) in combination with the *esg-QF* driver during diapause. Using this system makes it possible to monitor the cell cycle at cold temperatures, and we found that a significantly higher proportion of *esg*+ cells were in the G2/M phase during diapause compared to non-diapause controls (Fig. 2K, 2L). Additionally, using the *CycB-GFP* line, we confirmed increased *CycB* expression in ISCs during diapause (Fig. S2H), reinforcing the idea that ISCs arrest in the G2/M phase. These results suggest that ISCs undergo cell cycle arrest in the G2/M phase during dormancy.

Combined, our findings reveal that ISCs undergo drastic behavioral changes during DDMR–that they effectively halt the cell cycle at a recoverable state, reducing the number of cells produced post-eclosion and allowing for proper size change during diapause.

### Molecular signatures of DDMR

To explore the molecular signature of DDMR, we conducted a transcriptome analysis of the midgut. Specifically, bulk RNA sequencing (RNA-seq) was performed on samples of the entire adult midgut from two conditions: 3 weeks diapause (3wD) and 3 weeks non-diapause control (3wC). We identified 3,391 genes that were differentially expressed during dormancy (Fig. 3A). Gene Ontology (GO) enrichment analysis and Reactome pathway analyses revealed that metabolic processes, including small molecule, lipid, and carboxylic acid metabolism were significantly enriched in the 3wD midgut (Fig. S3A, S3B), suggesting that large-scale metabolic remodeling occurs alongside structural changes in the midgut during dormancy. Lipid metabolism-related genes such as *bmm* and *pepck*, along with carbohydrate metabolism-related genes like *tobi*, were differently expressed (Fig. S3C). Additionally, pathways related to metabolism (e.g., Metabolism of Lipids), translation regulation (e.g., Translation), and circadian gene degradation (e.g., Degradation of TIM) were among the top processes identified (Fig. S3B). Notably, circadian genes *per* and *tim*, which are upregulated in the entire body during diapause (Kučerová et al., 2016), also showed significantly increased expression in the midgut during dormancy (Fig. S3D). Furthermore, the expression of the antimicrobial peptide *Drs*, downstream of the Toll pathway, dramatically increased during dormancy (Fig. S3E), reflecting enhanced innate immunity, a hallmark of diapause at the organismal level (Kubrak et al., 2014).

**Figure 3.**
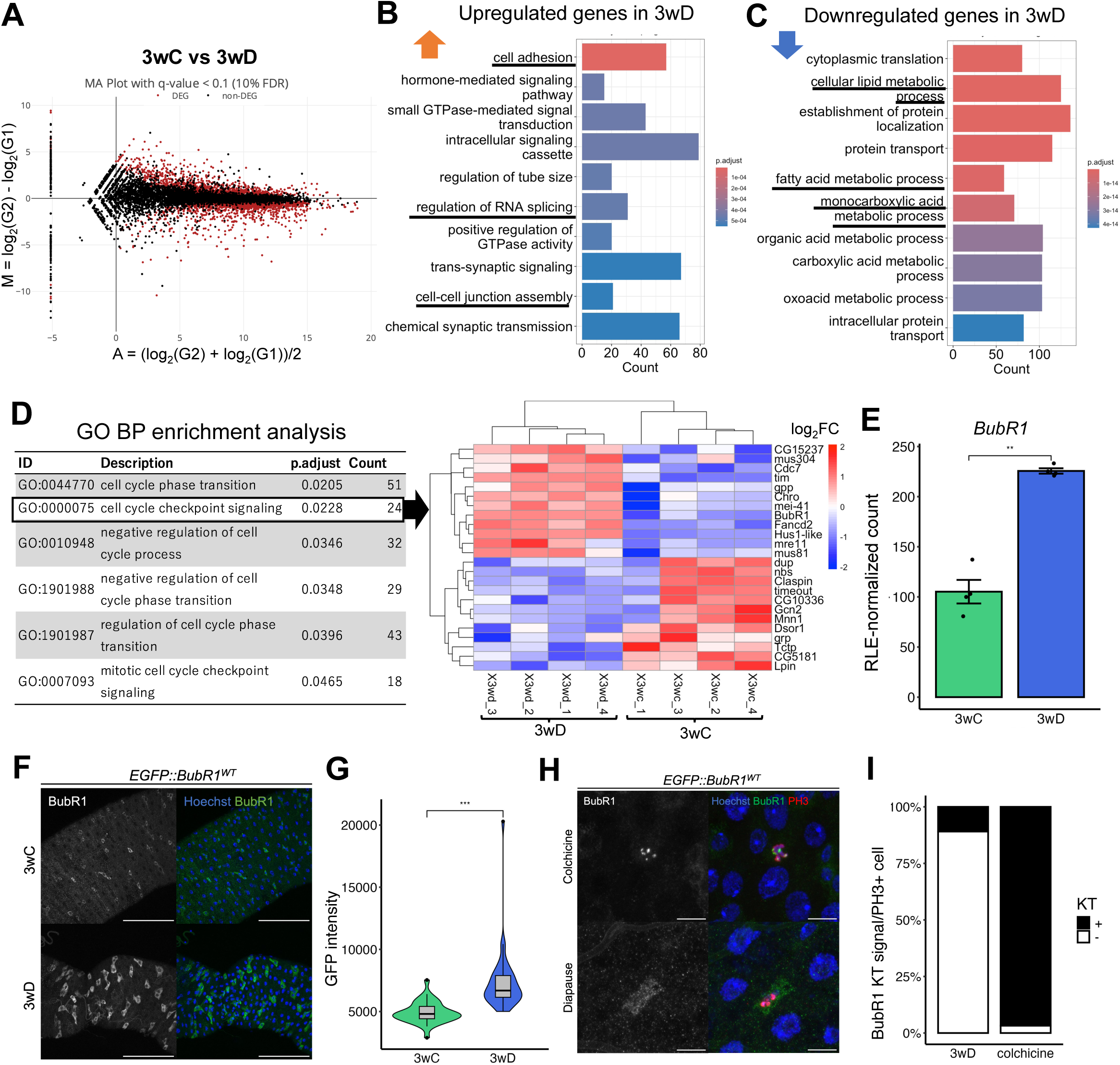
BubR1 expression increases in the cytoplasm of ISCs during diapause. (A) MA plot of differentially expressed genes in a pair-wise comparison of 3wC and 3wD conditions. (B, C) Gene ontology (GO) enrichment for upregulated genes (B) and downregulated genes (C) in 3wD. (D) Cell-cycle-related GO terms (biological process, BP) enriched in 3wD (left) and heat map of “cell cycle checkpoint signaling” genes (right). These GO terms were identified through enrichment analysis of all differentially expressed genes. (E) Expression level of *BubR1* in 3wC and 3wD midguts from RNA-seq data. Count data was normalized with RLE (Relative Log Expression). *n*=4 (3wC), 4 (3wD) RNA samples. (F) Representative images of EGFP::BubR1 in the midgut under 3wC and 3wD conditions. Scale bars: 50 μm. (G) Quantification of the EGFP::BubR1 signal intensity of (F). 30 cells from 4 midguts (3wC), 41 cells from 4 midguts (3wD). (H) Representative images of BubR1 localization in ISCs. BubR1 is localized in the cytosol under the 3wD condition while BubR1 is accumulated at the kinetochore (KT) after colchicine treatment. Scale bars: 10 μm. (I) The proportion of BubR1 localization in mitotic (PH3+) cells. 215 cells from 6 midguts (3wD), 31 cells from 4 midguts (Colchicine fed fly).

A detailed enrichment analysis was performed on the differentially expressed genes, by categorizing the genes into two groups: those upregulated during dormancy and those downregulated. In the upregulated gene group, GO terms associated with cell adhesion and RNA processing (e.g., regulation of RNA splicing) were identified as top-ranked categories (Fig. 3B). Genes associated with RNA processing and RNA metabolism are also upregulated in the ovaries during diapause, suggesting the existence of dormancy-specific translational regulation conserved across organs (DiVito Evans et al., 2023). In the downregulated gene group, by contrast, GO terms related to metabolic processes (e.g., cellular lipid metabolic process) were prominently enriched (Fig 3C). During diapause, extensive changes in the expression of genes involved in metabolic pathways have been observed throughout the body (Kubrak et al., 2014; Kučerová et al., 2016), and these findings suggest that gene expression changes linked to DDMR globally follow a pattern similar to that observed in the entire organism and other organs.

Next, to explore the molecular signature of DDMR recovery, RNA-seq was performed using samples of the entire adult midgut from two conditions: 3 weeks diapause followed by recovery (3wD+R) and 3 weeks control followed by recovery (3wC+R). We identified 526 genes with significant expression changes between the 3wD+R and 3wC+R midguts (Fig. S3G). GO enrichment and Reactome pathway analyses identified significant enrichment of genes associated with muscle differentiation and smooth muscle contraction (Fig. S3H, S3I). Given the extensive structural remodeling of muscles during insect dormancy (Kim et al., 2006; Kim & Denlinger, 2009), these gene expression changes related to muscle structure during recovery may contribute to the resizing of the midgut. Consistent with this finding, *dilp3*, which codes the product secreted by muscles during feeding and promote ISC proliferation (O’Brien et al., 2011), was significantly upregulated in the 3wD+R midgut (Fig. S3J), suggesting growth-promoting interactions between muscle and ISCs during recovery. In addition, neuropeptide genes such as *NPF, Dh31,* and *CCHa2* showed significant increases in expression (Fig. S3J), indicating that hormone secretion from EEs may change during recovery, potentially influencing gut growth and inter-organ communications. Overall, our findings suggest that the midgut undergoes large-scale structural and metabolic remodeling associated with extensive gene expression changes.

### BubR1 expression increases in the cytoplasm of ISCs during diapause

Because ISC cell cycle are arrested during diapause (Fig. 2), we focused on differentially expressed genes categorized under the GO term “cell cycle checkpoint signaling” (Fig. 3D). G2/M phase checkpoint regulators such as *mre11*, *mei-41* showed significant changes while *CG15237* and *BubR1* were differentially expressed as M-phase checkpoint regulators. These gene expression changes align with ISC behaviors during dormancy such as prolonged mitotic phases and an increase in G2/M phase cells. Among these genes, *BubR1* is particularly noteworthy due to its established roles as a mitotic checkpoint regulator and mitosis timer (Karess et al., 2013) and its expression in ISCs (Shinoda et al., 2023). Importantly, *BubR1* expression increased nearly threefold during diapause (Fig. 3E), suggesting a major potential impact on ISC behaviors and midgut remodeling.

To investigate *BubR1* expression at the cellular level, we used fly lines, which allow for the detection of the endogenous expression pattern of *BubR1* (*EGFP::BubR1^WT^* and *GFP-BubR1*) to visualize its expression in midgut cells. During reproductive dormancy, *BubR1* was strongly expressed in the cytoplasm of ISCs, primarily in R4 regions (Fig. 3F, 3G, S3F). This cytoplasmic localization during diapause contrasts with the localization of BubR1 associated with typical spindle checkpoint during mitosis, where it accumulates around kinetochores (Resende et al., 2018). Indeed, colchicine treatment, which induces mitotic arrest through the spindle assembly checkpoint (SAC), caused strong BubR1 localization around kinetochores in ISCs under non-diapause conditions (Fig. 3H, 3I). In contrast, during diapause, despite ISCs remaining arrested in mitosis, BubR1 remained cytoplasmic, with minimal signal around the kinetochores (Fig. 3H, 3I). These findings suggest that the role of BubR1 during DDMR is likely spindle checkpoint-independent and constitutes a unique function in the midgut during the dormancy state.

### BubR1 expression in ISCs is essential for DDMR

To explore the function of BubR1 in reproductive dormancy, we examined its effects on midgut remodeling using two *BubR1* mutant alleles *BubR1^k06109^* and *BubR1^k03113^*. qPCR analysis revealed that midgut *BubR1* expression in diapause-induced flies was significantly reduced in both heterozygous mutants, with *BubR1^k03113^*showing 48% and *BubR1^k06109^* showing 57% expression compared to controls (Fig. 4A). We then measured midgut size during diapause in these mutants and found a significant increase in midgut size compared to control flies (Fig. 4B, 4C). This midgut enlargement was accompanied by an increase in the total cell number, the number of mitotic ISCs, as well as the number of *esg+* cells (Fig. 4D-4F). Importantly, when BubR1 function was rescued in the *BubR1* trans-heterozygous mutant (*BubR1^k06109^*/*BubR1^k03113^)* with an *EGFP::BubR1^WT^*transgene (Shinoda et al., 2023), the total cell number in the posterior region and mitotic cell number in the midgut returned to levels comparable to controls (Fig. 4C-4E). These results strongly suggest that BubR1 is required for proper midgut remodeling during diapause. Notably, no ovarian development was observed in any of the dormant mutants, confirming the successful induction of ovarian dormancy despite impaired midgut remodeling (data not shown). Furthermore, no significant differences in midgut size were detected between these mutant backgrounds under non-dormant conditions (Fig. S4A), consistent with the result that the number of mitotic cells did not increase—or even decreased— in *BubR1* mutants (Fig. S4B). Collectively, these results highlight a critical role for BubR1 in DDMR.

**Figure 4.**
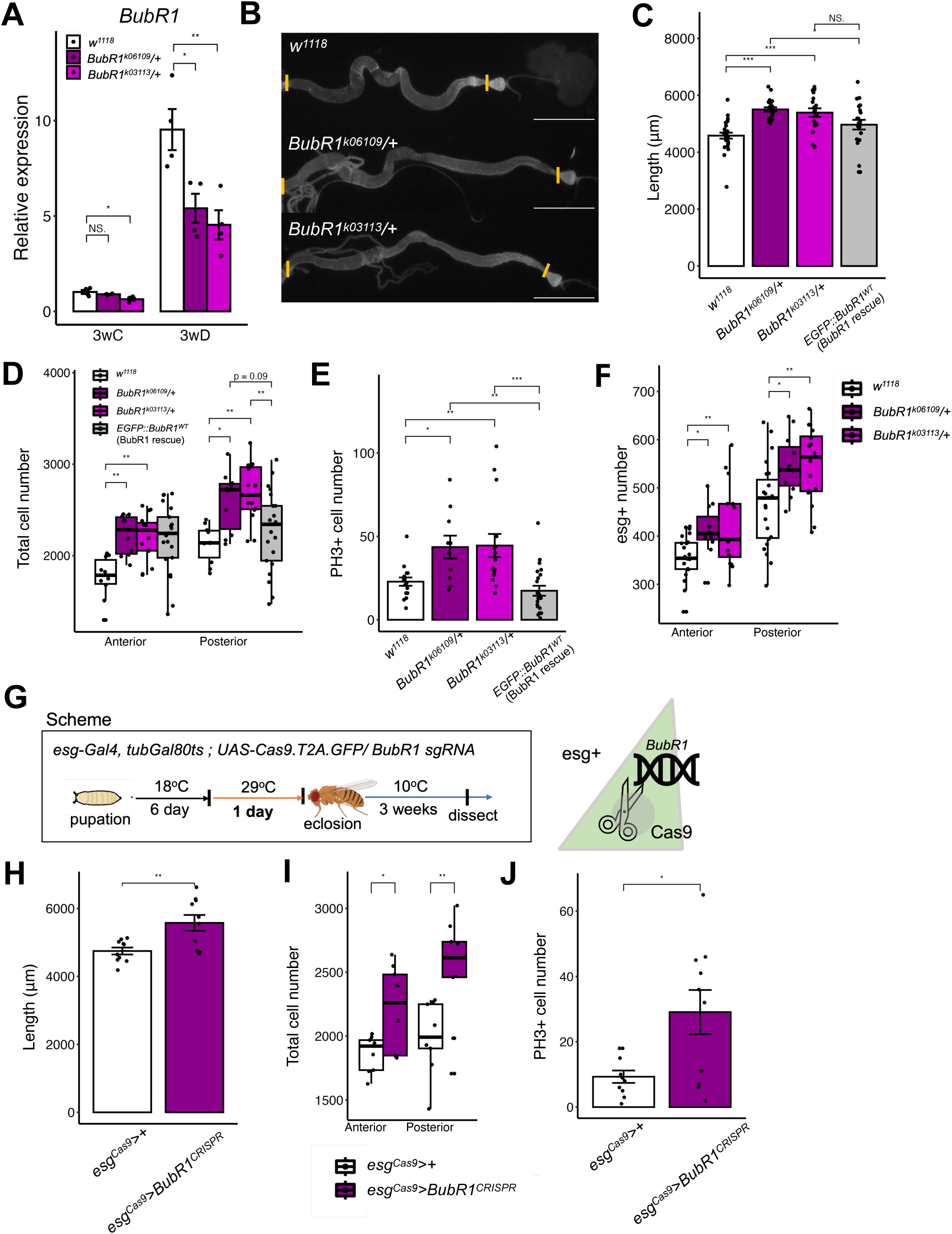
BubR1 expression in ISCs is essential for DDMR. (A) RT-qPCR of *BubR1* in control and *BubR1* heterozygous mutant flies under the 3wC and 3wD condition. *n*=4 (3wC, *w^1118^*), 3 (3wC, *BubR1^k06109^/+*), 4 (3wC, *BubR1^k03113^/+*), 4 (3wD, *w^1118^*), 4 (3wD, *BubR1^k06109^/+*), 4 (3wD, *BubR1^k03113^/+*) RNA samples. (B) Representative images of the midgut in control and *BubR1* heterozygous mutant flies under the 3wD conditions. Scale bars: 1 mm. (C) Quantification of the midgut length in control, *BubR1* heterozygous mutant, and *EGFP::BubR1^WT^* flies under 3wD. In *EGFP::BubR1^WT^* flies, wild-type form of *BubR1* is BAC rescued in the *BubR1* trans-heterozygous mutant background. *n*=27 (*w^1118^*), 22 (*BubR1^k06109^/+*), 18 (*BubR1^k03113^/+*), 23 (*EGFP::BubR1^WT^*) midguts. (D) Total cell number in control, *BubR1* heterozygous mutants, and *EGFP::BubR1^WT^*rescue flies under 3wD. *n*=10 (*w^1118^*), 10 (*BubR1^k06109^/+*), 15 (*BubR1^k03113^/+*), 23 (*EGFP::BubR1^WT^*) midguts. (E) PH3+ cell number in control, *BubR1* heterozygous mutants, and *EGFP::BubR1^WT^*rescue flies under 3wD. *n*=16 (*w^1118^*), 10 (*BubR1^k06109^/+*), 15 (*BubR1^k03113^/+*), 22 (*EGFP::BubR1^WT^*) midguts. (F) Quantification of *esg*+ cell number in control, *BubR1* heterozygous mutants under 3wD. *n*=23 (*w^1118^*), 11 (*BubR1^k06109^/+*), 16 (*BubR1^k03113^/+*) midguts. (G) Schematic of CRISPR-mediated *BubR1* knockout in *esg*+ cells. (H) Quantification of midgut length in control and *esg*+-specific *BubR1* knockout flies. *n*=10 (*esg^Cas9^>+*),10 (*esg^Cas9^>BubR1^CRISPR^*) midguts. (I) Total cell number in control and *esg*+-specific *BubR1* knockout flies. *n*=9 (*esg^Cas9^>+*),9 (*esg^Cas9^>BubR1^CRISPR^*) midguts. (J) PH3+ cell number in control and *esg*+-specific *BubR1* knockout flies. *n*=10 (*esg^Cas9^>+*),10 (*esg^Cas9^>BubR1^CRISPR^*) midguts.

The results obtained from the mutants could potentially reflect non-autonomous effects of *BubR1* expressed in other tissues. We thus investigated whether DDMR is influenced by BubR1 activity specifically in adult ISCs. To this end, we employed somatic tissue-specific CRISPR-mediated depletion (Bier et al., 2018; Meltzer et al., 2019), combined with the temperature-sensitive Gal80^ts^ system (McGuire et al., 2003) to induce *BubR1* mutations in ISC lineages before eclosion (Fig. 4G, S4C). When adult flies were shifted to dormancy conditions, we observed a significant increase in midgut size in ISC/EB-specific *BubR1* knockout flies (Fig. 4H). Furthermore, the number of total and mitotic cells also increased in these somatic *BubR1* knockout lines (Fig. 4I, 4J). Altogether, our results demonstrate that *BubR1* function in ISCs is essential for midgut remodeling during reproductive diapause.

Intriguingly, in *D. bifasciata* and *D. triauraria,* which undergo a programmed state of diapause in response to short-day length, *BubR1* expression in the midgut is higher in diapausing flies under SD compared to non-diapausing flies under LD (Fig. S4D, S4E). These results suggest a potential role of BubR1 in DDMR for the genus *Drosophila* more broadly.

### The MCC component Mad2 is involved in DDMR

During mitosis, BubR1 forms a complex with Mad2, Bub3, and Cdc20, which constitute key components of the mitotic checkpoint complex (MCC) and function cooperatively at the kinetochores (Musacchio, 2015). Our RNA-seq analysis indicated that, in addition to *BubR1,* the expression levels of *mad2* increased significantly during dormancy (Fig. 5A), though it was not identified as a differentially expressed gene. Intriguingly, Mad2 can interact with BubR1 in the cytoplasm of mammalian cells to regulate insulin signaling and metabolism (Choi et al., 2016). This raises the possibility that Mad2 may play a critical role in DDMR, independently of its role in the MCC.

**Figure 5.**
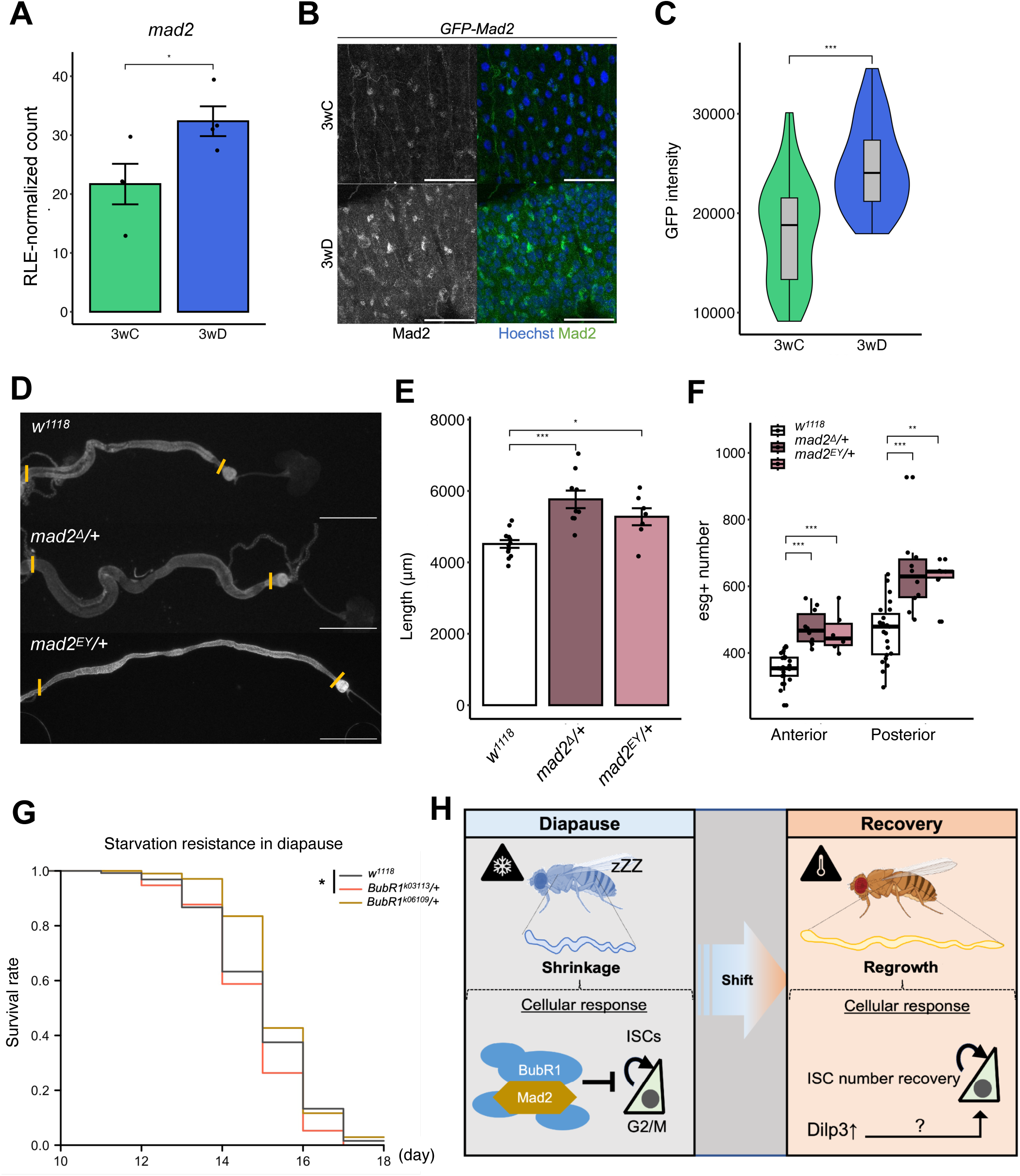
The MCC component Mad2 is involved in DDMR. (A) Expression level of *mad2* in 3wC and 3wD midguts from RNA-seq data. Count data was normalized with RLE. (B) Representative images of *GFP-Mad2* midguts under 3wC and 3wD conditions. (C) Quantification of GFP-Mad2 signal intensity of (B). 40 cells from 4 midguts (3wC), 52 cells from 4 midguts (3wD). (D) Representative images of the midgut in control and *mad2* heterozygous mutant flies under 3wD. Scale bars: 1 mm. (E) Quantification of midgut length in control and *mad2* heterozygous mutant flies under 3wD. *n*=12 (*w^1118^*), 9 (*mad2^Δ^/+*), 7 (*mad2^EY^/+*) midguts. The same *w^1118^* control samples were used in Figure 5E and Figure 4C. (F) Quantification of *esg*+ cell number in control and *mad2* heterozygous mutant flies under 3wD. *n*=23 (*w^1118^*), 10 (*mad2^Δ^/+*), 6 (*mad2^EY^/+*) midguts. The *w^1118^*control are same sample as in Figure 4E. (G) Starvation resistance in control and *BubR1* heterozygous mutant flies. *n*=128 (*w^1118^*), 114 (*BubR1^k06109^/+*), 103 (*BubR1^k03113^/+*) flies. (H) Graphical summary of midgut remodeling in diapause and recovery conditions.

To assess this potential role, we utilized the *GFP-Mad2* line to examine *mad2* expression in the adult midgut and found it to be highly expressed in the cytoplasm of ISCs during diapause (Fig. 5B, 5C), reminiscent of the cytosolic localization of BubR1 (Fig. 3F, S3F). To further investigate the function of Mad2 during DDMR, we analyzed midgut size using two *mad2* mutant alleles: *mad2^Δ^*and *mad2^EY21687^*. Our analysis revealed that heterozygous *mad2* mutants had significantly larger midguts during dormancy compared to controls (Fig. 5D, 5E). We also found that the number of *esg*+ cells increased in the anterior and posterior region of the midgut (Fig. 5F). These findings suggest that during reproductive diapause, Mad2 is localized in the cytoplasm of ISCs similarly to BubR1 and that the two proteins together regulate DDMR.

### DDMR enhances starvation resistance

Halting organ growth during dormancy is thought to conserve energy, thereby enhancing survival during periods of food scarcity, such as winter (Denlinger, 2022). In *Drosophila*, maintaining midgut size during reproductive diapause may contribute to increased resistance to starvation. To test this hypothesis, we compared starvation resistance between dormant *BubR1* mutants and controls. Although dormancy generally enhances starvation resistance compared to non-dormant states, *BubR1 ^k03113^* mutant flies exhibited significantly reduced starvation resistance during dormancy (Fig. 5G). These results suggest that BubR1/Mad2-mediated DDMR plays a critical role in enhancing starvation resistance at the organismal level.

## Discussion

In this study, we demonstrate that BubR1 and Mad2 play crucial roles in DDMR in *Drosophila* by modulating ISC dynamics to achieve diapause-specific states. While BubR1 and Mad2 are well-known as components of the MCC that localizes to the kinetochore, we found that in the diapausing midgut, they are predominantly localized in the cytoplasm of ISCs. Depletion of these proteins causes misregulated midgut growth and leads to reduced starvation resistance during diapause. Based on these findings, we propose that BubR1 and Mad2 act as key regulators of the ISC cell cycle during diapause, enabling a reversible state that facilitates recovery (Fig. 5H).

The *Drosophila* midgut undergoes extensive remodeling in response to various internal and external stimuli that directly or indirectly influence ISC activity (Christensen et al., 2024; Nagai et al., 2022; Stojanović et al., 2022). During starvation, for instance, the midgut size decreases as ECs undergo apoptosis, while upon refeeding, ISCs are stimulated by Dilp3 secreted from visceral muscles, promoting symmetrical ISC division and restoring midgut size (O’Brien et al., 2011). Similarly, during reproductive diapause—wherein food intake is reduced by less than half (Kubrak et al., 2014)—mild starvation responses may also contribute to midgut size reduction. Indeed, reproductive arrest can be more readily induced by adding starvation conditions (Ojima et al., 2018). Our RNA-seq data revealed increased *dilp3* expression and decreased expression of the FoxO target gene *Thor* during recovery (Fig. S3J), suggesting the possibility that Dilp3 secreted from muscles contributes to ISC activation as it does in post-starvation refeeding. However, in contrast to the simple starvation response, midgut cell death is minimal during diapause (Fig.S2I), and ISC cell cycles are arrested at G2/M in a recoverable state (Fig. 2). Furthermore, during reproductive arrest, germline stem cells in ovaries exhibit behaviors markedly different from those observed during amino acid deficiency responses (Easwaran et al., 2022). Collectively, these findings suggest that organ-level responses during diapause are driven by a complex interplay of environmental factors rather than by a simple starvation-induced mechanism.

Across species, developmental diapause typically involves a temporary halt in the cell cycle that delays organ growth. For example, during pupal diapause in the *Sarcophaga crassipalpis* brain and embryonic diapause in the turquoise killifish, the cell cycle primarily arrests at the G0/G1 phase (Meller et al., 2012; Tammariello & Denlinger, 1998). By contrast, most cells in silkworm embryos are arrested in the G2 phase during diapause (Nakagaki et al., 1991). In this study, we observed a significant increase in ISCs arrested at the G2/M phase during DDMR. *Drosophila* oogenesis arrests during diapause due to activation of the Chk2 DNA damage checkpoint (Easwaran et al., 2022), and our findings suggest that similar mechanisms may govern growth arrest in the midgut. Indeed, genes like *mre11* that are differentially expressed in the diapausing midgut (Fig. 3D) are required for G2/M checkpoint activation (Song, 2005). Notably, ISCs exhibit active cell cycle arrest during diapause via checkpoint mechanisms, rather than undergoing a passive delay induced by low temperatures (Fig. S1E-S1F). This controlled cell cycle arrest in resident stem cells may contribute to the pausing of organ growth in other diapausing or hibernating animals and to facilitating rapid recovery once stem cell division resumes.

BubR1 is a highly conserved eukaryotic protein that functions as a key component of the SAC, forming the MCC with Mad2, Bub3, and Cdc20 to prevent premature progression of cell division and ensure accurate chromosomal segregation, and its disruption increases the risk of aneuploidy (Baker et al., 2013; Karess et al., 2013; Resende et al., 2018). Beyond these canonical functions, studies across cellular and organismal contexts have uncovered additional roles for BubR1. For example, in mouse embryonic fibroblasts, *BubR1* heterozygosity reduces *p53* and *p21* expression after DNA damage, impairing proper G2/M phase arrest (Fang et al., 2006). Additionally, BubR1, together with Mad2, recruits adaptor protein 2 to the insulin receptor (IR) to promote IR endocytosis and reduce insulin sensitivity (Choi et al., 2016; Park et al., 2023; Tang et al., 2020). Consistent with these findings, *BubR1* deficiency in mice leads to premature aging and reduced lifespan, highlighting its role in organism-wide responses (Baker et al., 2004). In *Drosophila*, BubR1 in the fat body negatively regulates *bmm* expression via IMD signaling during starvation, suppressing lipid breakdown and enhancing lifespan (Liu et al., 2024). These findings delineate a complex regulatory network involving BubR1 and associated proteins across different biological contexts. Although the regulation and interacting partners of BubR1 and Mad2 during diapause have yet to be identified, the observed changes in their expression in ISCs likely contribute to organism-level responses, including enhanced starvation resistance mediated by structural and metabolic changes in the midgut. Future investigations into these emerging additional roles of BubR1/Mad2 will enhance our understanding of organ remodeling in dormancy programs and provide new insights into the control of resident stem cells.

## Acknowledgements

We thank R. Giet, B. Ohlstein, C. Sunkel, N. Shinoda, BDSC, Kyoto Stock Center, NIG-FLY, KYORIN-Fly, and Developmental Studies Hybridoma Bank (DSHB) for fly stocks and reagents; Y. Hara and S. Kondo for discussion.

## Author contributions

Conceptualization: Y.A., Y.N.

Investigation: Y.A., H.N., Y.N.

Methodology: Y.A., H.N., T. F., A.T., Y.N.

Validation: Y.A., H.N., M.M., T. F., A.T., Y.N.

Data curation: Y.A., Y.N.

Writing – original draft: Y.A., H.N., Y.N.

Writing – review & editing: Y.A., H.N., A.T., M.M., Y.N.

Supervision: M.M., Y.N.

Funding acquisition: H.N., M.M., Y.N.

## Materials and methods

### *Drosophila* stock and maintenance

All stocks were maintained on a standard diet containing 4% cornmeal, 6% baker’s yeast (Saf Yeast), 6% glucose (Wako, 049-31177), and 0.8% agar (Kishida chemical, 260-01705) with 0.3% propionic acid (Tokyo Chemical Industry, P0500) and 0.05% nipagin (Wako, 132-02635).

Canton-S was utilized as the wild-type strain. Transgenic fly lines were obtained from the Bloomington Drosophila Stock Center (BDSC), NIG-Fly and the Kyoto Stock Center. Unless otherwise indicated, strain descriptions can be found at Flybase (http://flybase.bio.indiana.edu): *Dl-lacZ* (Röttgen et al., 1998) (BDSC 11651), *esg-QF2* (Nagai et al., 2023)*, QUAS-Fly-FUCCI* (Zielke et al., 2014) (BDSC 55108), *BubR1^k03113^* (Kyoto 102258), *BubR1^k06109^* (Kyoto 102444), *mad2^EY^* (Bellen et al., 2011) (BDSC 22495), *tub-Gal80^ts^* (BDSC 7017, 7019, described in Flybase and BDSC), *esg-Gal4* (Goto & Hayashi, 1999) (Kyoto 109126), *UAS-GFP* (BDSC 1521), *UAS-Cas9.T2A.eGFP* (Trivedi et al., 2020) (BDSC 94298), *FRT40A* (originated from BDSC 5074), *attP40{U6-gRNA}* (NIG 2RG-1578). The following lines were gifts from fly community: *esg-GFP[P01986]* (Le Bras & Van Doren, 2006), *Su(H)GBE-lacZ* (Furriols & Bray, 2001), *yw hs-flp; tub-Gal80 FRT40A; tub-Gal4 UAS-dsRed /TM6B* (Guo et al., 2013)*, EGFP::BubR1^WT^* (Shinoda et al., 2023)*, GFP-BubR1, GFP-Mad2* (Buffin et al., 2005)*, mad2^Δ^* (Buffin et al., 2007)*, CycB-GFP* (Huang & Raff, 1999). The following other *Drosophila* species were obtained from KYORIN-Fly: *D. bifasciata* (JZK3), *D. triauraria* (TM15-92), *D. triauraria* (KMJ1), *D. triauraria* (OEB12). See Additional file: Table S1 for the genotypes in each figure. We used female flies unless otherwise noted in the figures.

### Diapause and recovery induction

To assess the effects of diapause on midgut growth using *D. melanogaster*, we subjected old virgin females less than 6 h to 10°C and 8 L:16D photoperiods (Diapause condition), and 18°C at 12L:12D photoperiods (Control condition) for 1-5 weeks. We then allowed the flies to recover by transferring them into vials with fresh food with dry yeast for 7 days at 25°C and 12L:12D photoperiods (Recovery condition).

To assess the effects of the cold on midgut size, we allowed the flies to grow in 18°C for 4 days after eclosion, and then subjected them to 10°C and 8 L:16D photoperiods for 3 weeks (Cold condition).

To assess the effects of diapause state on midgut growth in other *Drosophila* species, we subjected virgin females less than 6 h old to 10 L:14D photoperiods (SD, diapause) and 16L:8D photoperiods (LD, control) for 16 days. The rearing temperature was kept consistent at 15°C. We then allowed the flies to recover by transferring them into 16L:8D photoperiod condition.

### MARCM clone induction scheme

For the mosaic analysis with a repressible cell marker (MARCM) analysis, MARCM-ready flies (*hsFLP; tub-Gal80,FRT40A/FRT40A; tub-Gal4,UAS-dsRed*) were cultured at 25°C throughout pupal development, except during heat shock induction. When female larvae began metamorphosis, white pupae were placed into an empty vial at 25°C and designated as being at 0 h after puparium formation (APF). MARCM labels one of the two daughter cells after mitotic division. To label dividing ISCs, MARCM-ready flies were heat shocked at 48 h APF for 1 h (the time during which ISCs undergo asymmetric divisions (Guo & Ohlstein, 2015)). After eclosion, the size of the growing ISC clones was observed with anti-Pros immunofluorescence staining under 1 week diapause or 1 week control conditions. EE-only clones, which can be induced by labeling daughter EE during asymmetric division were excluded by immunostaining against the EE marker Pros, and the size of clones containing two or more ISCs was quantified.

### Tissue-specific somatic mutation of BubR1

Transgenic flies expressing BubR1-sgRNA (GGTGACGCCAGTTACTATTG) obtained from NIG gRNA stock collections were crossed to flies expressing *esg^ts^>Cas9* (*esg-Gal4 tubGal80^ts^ / CyO; UAS-Cas9 / TM3, Act-GFP*). Experimental crosses were maintained at 18°C to utilize Gal80^ts^ for suppressing Gal4 activity. When female larvae began metamorphosis, white pupae were placed into an empty vial at 18°C. After 6 days, when ISC division was completed, pupae were transferred to 29°C and reared for 1 day to introduce somatic mutations. These pupae were then transferred back to 18°C and reared under diapause or control conditions for 3 weeks after eclosion.

### Immunofluorescence

Adult midgut samples were dissected in 1 × PBS and fixed in 4% PFA for 30–45 min at room temperature (RT). The following primary antibodies were used with indicated dilution into 1 × PBS containing 0.5% BSA and 0.1% Triton X-100: rabbit anti-PH3 (Millipore 06–570, 1:1000), mouse anti-Prospero (DSHB MR1A, 1:100), rat anti-GFP (Nacalai tesque 04404–26, 1:400), rabbit anti-cDcp1 (Cell Signaling Technology 9578, 1:200), chicken anti-β-galactosidase (Abcam ab9361, 1:500), and mouse anti-Delta (DSHB C594.9B, 1:100). After overnight incubation with primary antibodies at 4°C, samples were incubated with fluorescent secondary antibodies (Jackson ImmunoResearch and Invitrogen, 1:500) for 1 h at RT. Hoechst 33,342 (Invitrogen, final concentration: 10 μg/ml) was used to visualize DNA. Samples were mounted in Slowfade Diamond (ThermoFisher, S36963) and imaged with a Zeiss LSM880 confocal microscope.

### RT-qPCR

Total RNA was purified from 7-10 midguts using the ReliaPrep RNA Tissue Miniprep System (Promega). cDNA was made from 100 or 200 ng of RNA using PrimeScript RT Reagent Kit (TaKaRa). Quantitative PCR was performed using TB Green Premix Ex Taq II (TaKaRa) and the QuantStudio 6 Flex Real-Time PCR System (ThermoFisher). *RpL32* was used as an internal control. Primers for *D. bifasciata* and *D. triauraria* were designed based on the sequence homology to *D. melanogaster* transcripts. Primer sequences are listed in Table S2.

### Bulk RNA-seq

Total RNA was purified from 10 to 15 midguts using the ReliaPrep RNA Tissue Miniprep System (Promega). A NanoDrop (ThermoFisher) was used to measure the concentration of each RNA sample, and the integrity of RNA samples was checked using a Bioanalyzer (Agilent). A total of 300 ng RNA for each sample was used for cDNA library preparation and the subsequent sequencing was performed on an DNB sequencing system with single reads of 50 base pairs. The sequencing reads were aligned against the *Drosophila melanogaster* genome (ENSEMBL, BDGP6.22) using Salmon (Patro et al., 2017) for transcript quantification. The quantification data of each transcript was converted into gene-level counts using Tximport (Soneson et al., 2015). Differential expression analysis of two conditions/group (four biological replicates per condition) was performed using the DESeq2 R package (ver 3.20). Genes with an adjusted *P*-value < 0.1 were assigned as differentially expressed. For functional cluster analysis of DEGs, GO BP enrichment analysis and Reactome pathway analysis were performed with the ClusterProfiler R package.

### Colchicine treatment

0.2 mM Colchicine was dissolved in 5% (w/v) sucrose solution. Filter paper (Whatman 3MM) was soaked with 300 μL of reagents and placed into empty vials. Newly-eclosed flies were transferred into these vials and fed with 5% sucrose only (control) or colchicine solutions for 1 day.

### Starvation resistance

To assay the starvation resistance, flies were fed with agar only food. For the survival assay, dead flies (determined by immobility and a lack of response to tapping) were counted daily.

### Quantification of midgut size

The central line length and area of the midgut were quantified using the Fiji/ImageJ free hand tool on images obtained with a Zeiss AXIO Zoom.V16 stereomicroscope.

### Quantification of cell number

Cell number was quantified using Fiji/ImageJ. To count the total cell and *esg-GFP*^+^ cell number, Hoechst and GFP signal was processed as follows: (1) despeckle, (2) binarization, (3) watershed, (4) analyze particle. Cell counts for PH3^+^ cells, *Su(H)GBE-lacZ*^+^ cells, and Fly-Fucci*^+^* cells were manually counted using the cell counter function.

For MARCM clone size quantification, clones containing two or more ISCs were selected, and the number of cells in the clone was counted manually.

### Quantification of signal intensity

The signal intensity of reporter lines (*EGFP::BubR1^WT^, GFP-Mad2*) was quantified using Fiji/ImageJ. The signal areas were selected as ROIs using the polygon selection tool. The reporter intensity in each ROI was quantified using the Measure command.

### Statistics

Statistical analyses were performed using Excel and RStudio. Two-tailed *t* tests were used for comparisons between two groups. One-way ANOVAs with post hoc Tukey tests were performed when comparing three or more groups. Log-rank tests were used for survival curve comparison. Significance is indicated in the figures as follows: \**P* ≤ 0.05, \*\**P* ≤ 0.01, \*\*\**P* ≤ 0.001, not significant (N.S.): *P* > 0.05. Bar graphs show mean ± standard error. Boxplots show median (thick line in the box), first and third quartiles (bottom and top of the box), minimum value (lower whisker), and maximum value (upper whisker). Dots in bar graphs and boxplots indicate individual values. Violin plots portray the distribution of individual values.

## Figure legends

**Figure S1. Midgut remodeling in coordination with *Drosophila* reproductive diapause**

(A) Representative image of ovary growth arrest in *D. melanogaster* diapause. Scale bar: 1 mm.

(B) Schematic of control and recovery in *D. melanogaster*. LD = Light: Dark.

(C) Quantification of midgut length of wild-type flies (Canton-S) under control and recovery conditions. *n=*6-13 midguts for each condition. The control samples are same as in Figure 1C.

(D) Quantification of midgut length in the recovery condition for 1-7 days. *n*=8-21 midguts for each condition.

(E) Schematic of the diapause and cold conditions in *D. melanogaster*. LD = Light: Dark. To set up the cold condition, we allowed the flies to grow in 18°C for 4 days after eclosion, and then held them at 10 °C with 8 L:16D photoperiods for 3 weeks.

(F) Quantification of midgut length under diapause and cold conditions after 3 weeks. *n*=9-21 midguts for each condition. The 3 weeks diapause samples are the same shown in Figure 1C.

(G) Quantification of midgut length and area in wild-type male flies (Canton-S) under 3wC and 3wD condition. *n*=7 (3wC), 7 (3wD) midguts.

(H) Representative images of ovaries in *D. bifasciata* (JZK3) and *D. triauraria* (TM15-92) under long day (LD) and short day (SD) conditions. In SD, ovarian growth arrest was induced. Scale bars: 1 mm.

(I, J) Quantification of midgut length of non-diapause strains of *D. triauraria* (KMJ1 and OEB12) under LD and SD conditions. *n*=11 (KMJ1, LD), 11 (KMJ1, SD), 8 (OEB12, LD), 13 (OEB12, SD) midguts. KMJ1 and OEB12 female flies do not enter diapause state in the SD condition (Yamada & Yamamoto, 2011).

**Figure S2. ISCs remain in a non-proliferative state during diapause**

(A) The proportion of esg+ cells in total cells. *n*=11 (3wC), 12 (3wD), 10 (3wD+R) midguts.

(B) Representative images of the posterior midgut under 3wC and 3wD conditions in *Dl-lacZ* flies. Scale bars: 100 μm.

(C) Representative images of anti-Dl staining in 3wC and 3wD wild-type flies. Scale bars: 100 μm.

(D) Representative images of the posterior midgut under 3wC and 3wD conditions in *Su(H)GBE-lacZ* flies. Scale bars: 100 μm.

(E) Quantification of the Su(H)+ cell number. *n*=8 (3wC), 6 (3wD) midguts.

(F) The proportion of Su(H)+ cells in total cells. *n*=8 (3wC), 6 (3wD) midguts.

(G) PH3+ cell number in the midgut of 3wC, 3wD, and 3wD+R flies. n=6 (3wC), 8 (3wD), 6 (3wD+R) midguts. The 3wC and 3wD samples are the same shown in Figure 2F.

(H) Representative images of the posterior midgut under 3wC and 3wD conditions in *CycB-GFP* flies. Scale bars: 100 μm.

(I) Representative images of anti-cDcp1 staining in 3wC and 3wD wild-type flies. Scale bars: 50 μm.

**Figure S3. Molecular signatures identified by RNA-seq analysis in DDMR**

(A) Gene ontology (GO) enrichment of 3wD transcriptome compared to 3wC. All differentially expressed genes in 3wD are used for enrichment analysis.

(B) Reactome Pathway Enrichment in 3wD compared to 3wC. All differentially expressed genes in 3wD are used for enrichment analysis.

(C) Expression levels of *bmm, pepck, tobi* from RNA-seq data. *n*=4 (3wC), 4 (3wD) RNA samples. Count data was normalized with RLE.

(D) Expression levels of *per, tim* from RNA-seq data. *n*=4 (3wC), 4 (3wD) RNA samples. Count data was normalized with RLE.

(E) Expression levels of *Drs* from RNA-seq data. *n*=4 (3wC), 4 (3wD) RNA samples. Count data was normalized with RLE.

(F) Representative images of GFP-BubR1 from 3wC and 3wD posterior midguts. Scale bar: 100 µm.

(G) MA plot of differentially expressed genes in a pair-wise comparison of 3wC+R and 3wD+R conditions.

(H) GO enrichment in 3wD+R compared to 3wC+R. All differentially expressed genes in 3wD+R are used for enrichment analysis.

(I) Reactome Pathway Enrichment in 3wD+R compared to 3wC+R. All differentially expressed genes in 3wD+R are used for enrichment analysis.

(J) Expression levels of *CCHa2, Dh31, dilp3, NPF, Thor* from RNA-seq data. *n*=4 (3wC+R), 4 (3wD+R) RNA samples. Count data was normalized with RLE.

**Figure S4. BubR1 functions in the midgut in coordination with *Drosophila* reproductive diapause**

(A) Quantification of midgut length in control and *BubR1* heterozygous mutants under 3wC. *n*=10 (*w^1118^*), 8 (*BubR1^k06109^/+*), 8 (*BubR1^k03113^/+*) midguts.

(B) PH3+ cell number in control, *BubR1* heterozygous mutants, and *EGFP::BubR1^WT^*flies under 3wC. *n*=10 (*w^1118^*), 8 (*BubR1^k06109^/+*), 8 (*BubR1^k03113^/+*) midguts.

(C) RT-qPCR of *BubR1* in control and CRISPR-mediated *BubR1* knockout flies under the 3wD condition. *n*=9 (*esg^Cas9^>+*), 9 (*esg^Cas9^>BubR1^CRISPR^*) RNA samples.

(D) RT-qPCR of *BubR1* in *D. bifasciata* (JZK3) midgut under the control (LD) and diapause (SD) conditions. *n*=8 (LD), 9 (SD) RNA samples.(E) RT-qPCR of *BubR1* in *D. triauraria* (TM15-92) midgut under the control (LD) and diapause (SD) conditions. *n*=6 (LD), 8 (SD) RNA samples.

