## Supplementary figures and images for "BubR1 and Mad2 regulate adult midgut remodeling in *Drosophila* diapause"

### Supplementary Figures

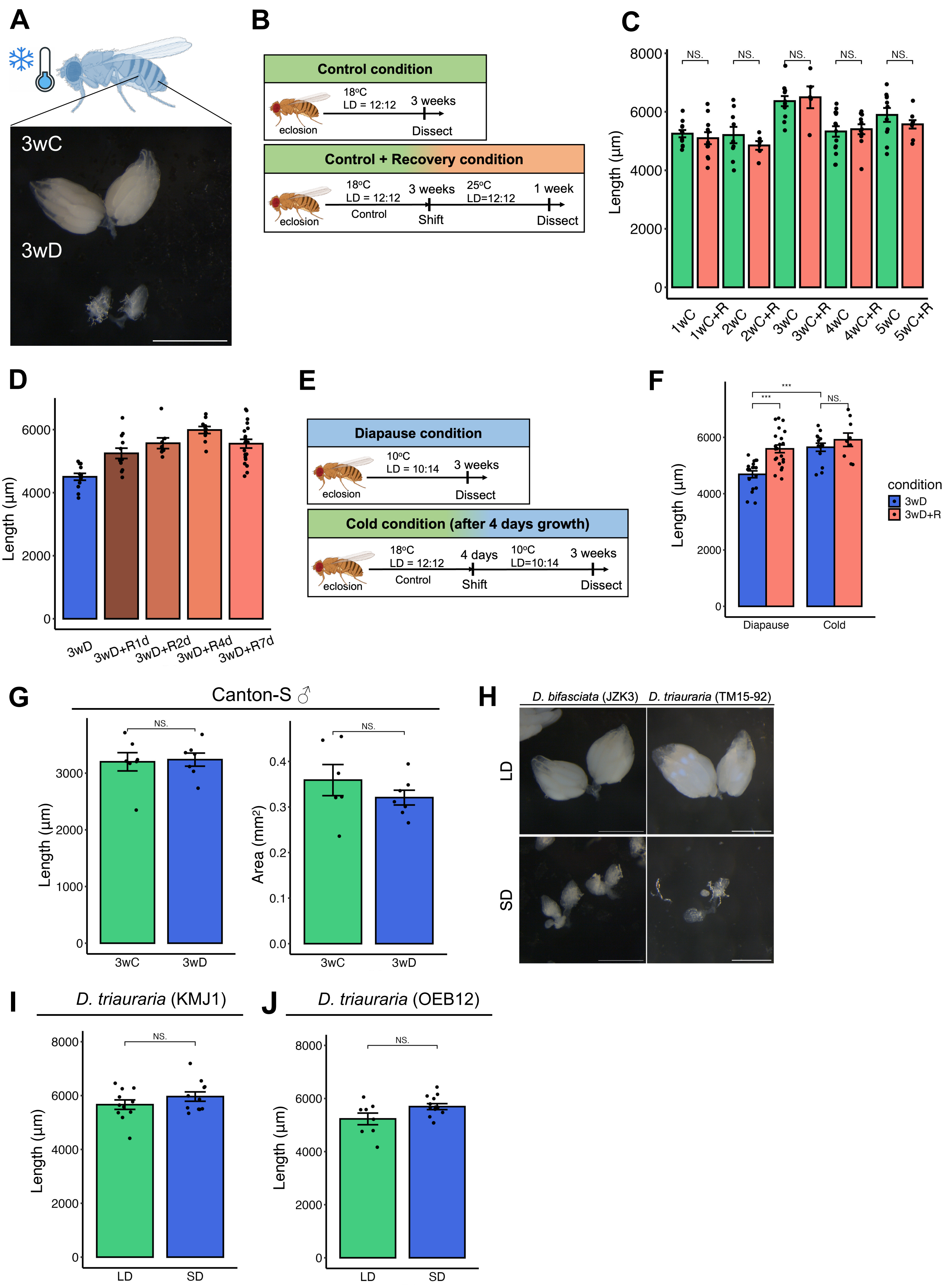

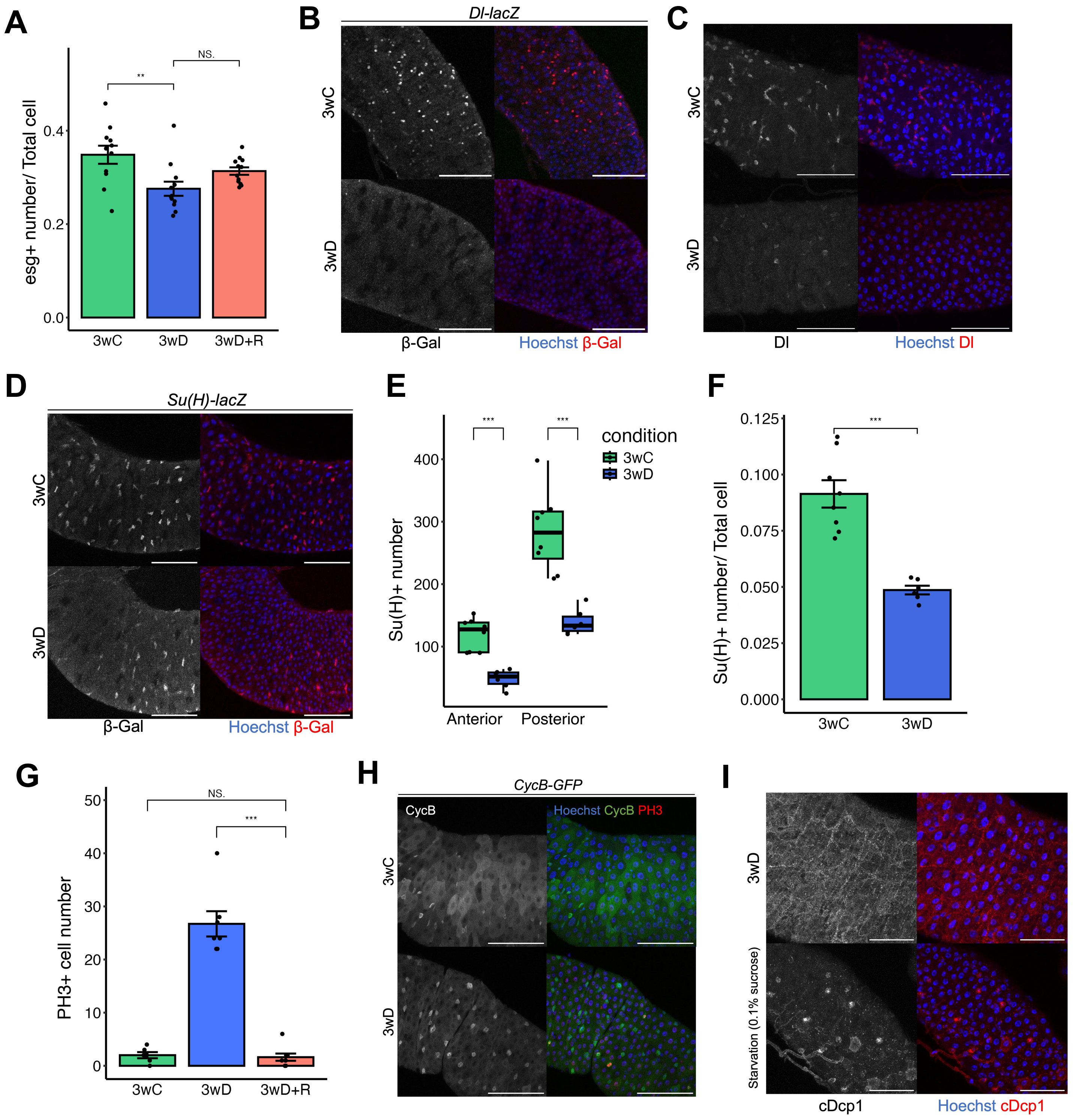

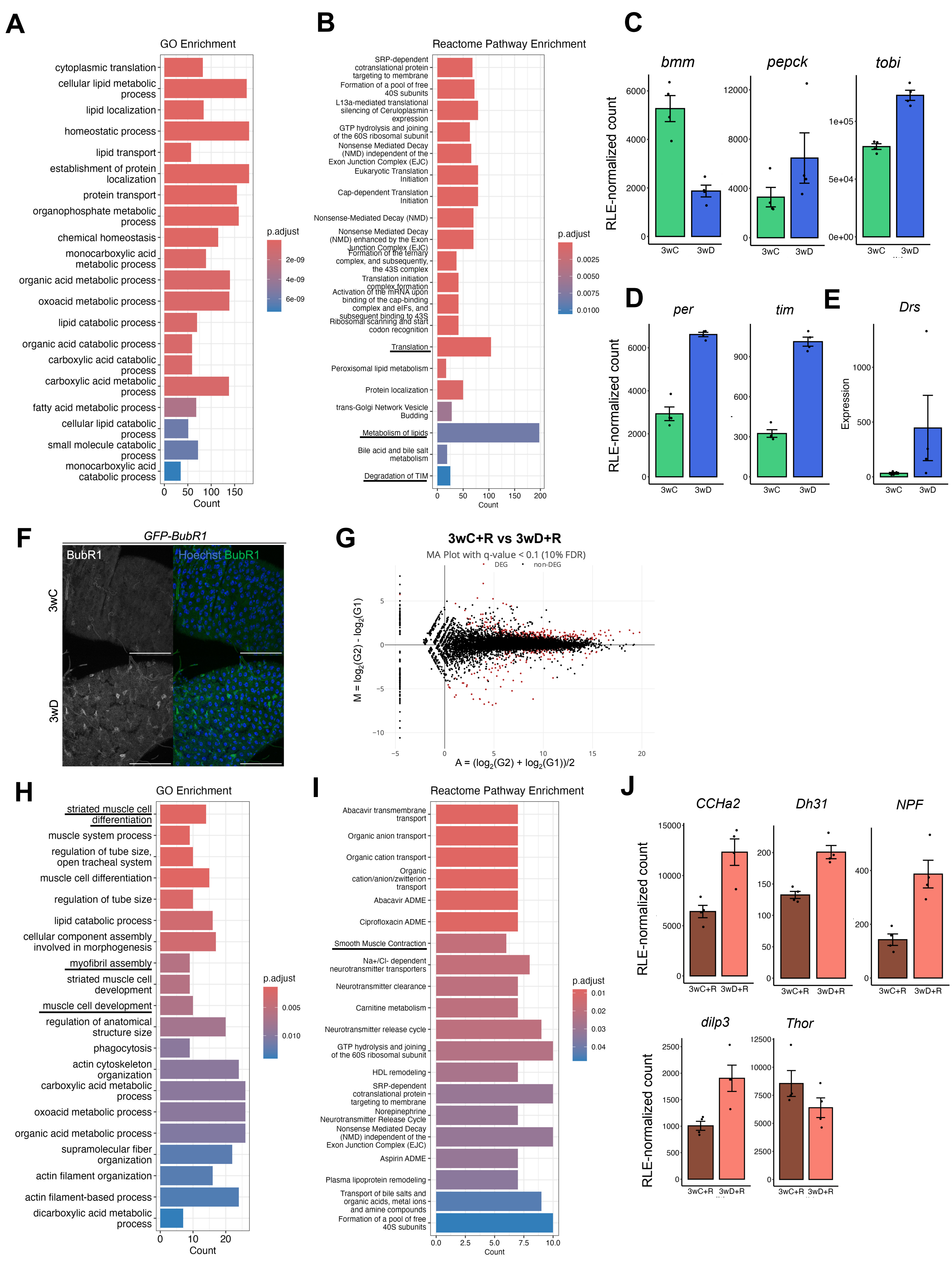

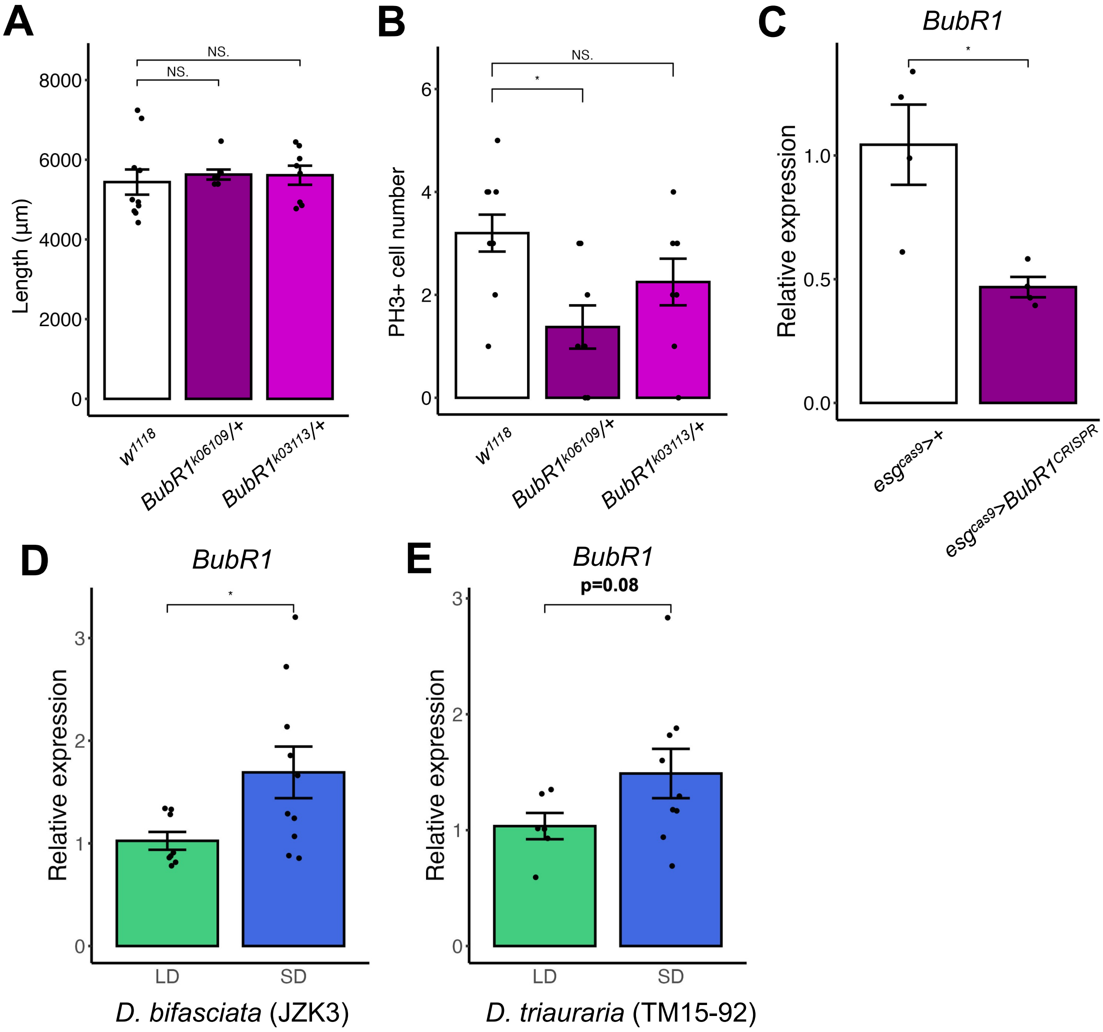
